# Auditory and visual mismatch negativity responses to numerical and non-numerical magnitudes in adults

**DOI:** 10.64898/2026.09.29.753775

**Authors:** Sadiq Mobeen, Tuire Koponen, Minna Torppa, Jarmo Hämäläinen, Kaisa Lohvansuu

**Affiliations:** Department of Psychology, University of Jyväskylä, Jyväskylä, Finland; Centre for Interdisciplinary Brain Research, University of Jyväskylä, Jyväskylä, Finland; Department of Education, University of Jyväskylä, Jyväskylä, Finland; Centre of Excellence for Learning Dynamics and Intervention Research (InterLearn), University of Jyväskylä, Jyväskylä, Finland; Department of Teacher Education, University of Jyväskylä, Jyväskylä, Finland

## Abstract

We investigated how auditory and visual presentation of small numerical magnitudes and non-numerical magnitudes are passively processed at the neural level. Earlier research suggests automatic processing of quantities occurs rapidly in the brain. However, there are no studies on passive, pre-attentive change detection of magnitudes utilizing mismatch negativity (MMN) across sensory modalities in same participants, important to understand amodal processing of magnitudes. We recorded EEG in passive auditory and visual multi-feature oddball and control paradigms from the same adult individuals (N=104/105, respectively). In the auditory paradigm, the standard stimulus was a 2-part sound, while the deviant stimuli varied in frequency, intensity, gap length, and number of sounds (1-part and 3-part stimuli). In the visual paradigm, the standard stimulus was a dot array containing two medium-sized dots, while the deviant stimuli varied in numerosity (1 or 3 dots) and dot size (small or large).

The results for the auditory paradigm showed significant MMN across all features, including comparisons with deviant-alone control stimuli, except for the 2-part vs. 1-part contrast. The results for the visual paradigm suggested differences in processing across all features apart from large and medium-size stimuli. These results replicate earlier MMN findings that the brain distinguishes pre-attentively changes in sound frequency, amplitude and gap. Auditorily presented small numerical magnitudes also showed pre-attentive processing but less robustly to the contrast 2 vs. 1. Visual pre-attentive processing of small numerical magnitudes was also observed. Together these findings could be interpreted to support the existence of abstract, pre-attentive processing of numerical magnitudes.

**Highlights:** • Auditory and visual processing of numerical and non-numerical magnitudes was examined with EEG

• Auditory non-numerical magnitudes and medium vs. small dots replicated earlier MMN studies

• aMMN and vMMN showed pre-attentive processing of changes in small numerosities 2 vs. 3 and 2 vs. 1

• 2 vs. 1 tone processing was less robust than 2 vs. 3 or the processing of the visual contrasts

• The brain processes changes in small non-symbolic magnitudes pre-attentively

## Introduction

Humans engage with non-symbolic and symbolic numerical information in various forms in their everyday lives, such as when examining temperature or speed, or when counting. Non-symbolic numerical cognition is thought to rely on two systems: the parallel individuation system for small numerosities and the approximate number system (ANS) for larger ones (Hyde, 2011). The behavioral manifestation of parallel individuation system for small numerosities, typically less than four, is referred to as subitizing (Kaufman et al., 1949).

According to recent evidence, ANS can activate even small numbers when attention is not directed to the stimuli whereas subitizing requires attentional involvement (Nieder, 2025). Additionally, numerical processing relies on the parallel individuation system in the visual domain. The parallel individuation system depends on the object tracking system (OTS) for small numbers of objects and likely requires attentional resources (Nieder, 2025; Pylyshyn, 2001). OTS emphasizes individuation, tracking and memory of items, and leads to mental representations of quantities (Hyde, 2011). Evidence suggests that small non-symbolic numerical magnitudes (e.g., dot arrays within the subitizing range) are processed by an OTS in the visual modality (Feigenson et al., 2004; Briggs et al., 2023) whereas in auditory modality (set of tones) the existence of a similar mechanism is yet to be explored within subitizing range. In this study, we focused on the passive, pre-attentive detection of changes in small non-symbolic numerosities in auditory and visual domains. We used auditory stimuli, sets of tones, varying in numerical and non-numerical magnitudes, and visual stimuli, sets of dots, to study how they are processed in the adult brain.

Processing magnitude in both auditory and visual modalities is suggested to originate in the intraparietal sulcus (IPS) (Dantzig, 2007; Dehaene, 1997). IPS is suggested to house an internal representation of numerical magnitude, making it a core region for numerical cognition especially in tasks involving symbolic and non-symbolic number comparison (Ansari et al., 2006). In addition to IPS, the prefrontal cortex has also been implicated in numerical processing. Neurons in the IPS activate earlier than those in the prefrontal cortex (PFC), indicating that numerical information may first be processed in the IPS before being relayed to the PFC for guiding number-related actions (Ansari, 2008). The horizontal segment of the intraparietal sulcus is engaged during numerical activities, highlighting its involvement in processing quantities regardless of how numbers are represented (Dehaene et al., 2003). In the IPS there are cells that respond to specific numbers, helping us quickly recognize and compare quantities (Santens et al., 2010; Fias et al., 2003). While the IPS is central to numerical processing, some researchers propose that numerical cognition recruits broader cognitive systems, including those involved in verbal and spatial processing (Van Opstal et al., 2012).

Processing of numerical and non-numerical magnitudes can be studied using EEG-based event-related potentials (ERPs). A particular ERP response, called the mismatch negativity (MMN), is useful for indexing pre-attentive change detection of deviant stimuli in a repeated stimulus stream. It was originally found in the auditory domain (Näätänen et al., 1978) but is present in visual processing and other modalities as well (Male, 2025). The brain’s automatic detection of change in visual stimuli elicits the visual MMN (vMMN) (Kimura et al., 2011; Stefanics et al., 2014; Male, 2025). vMMN has been consistently localized to parieto-occipital and occipital cortices, reflecting early sensory prediction error mechanisms (Male, 2025; Pazo-Alvarez et al., 2003). At parieto-occipital sites, vMMN typically emerges as a negative deflection occurring between 150 and 300 ms post-stimulus onset (Male, 2025).

However, one study showed that vMMN response can occur in an earlier time window within a 105–280 ms latency range in the same regions, suggesting temporal variability in cortical processing (File et al., 2017).

Auditory MMN (aMMN) has also been used in research to quantify objects and events during the pre-attentive discrimination of sounds with different auditory features such as frequency, duration, intensity, and gap duration (Näätänen et al., 2007). While aMMN has its main generators located in the auditory cortices (Woldorff et al., 1998; Näätänen et al., 2007), frontal regions, such as the inferior and middle frontal gyri, are also engaged during aMMN tasks, indicating their role in higher-order cognitive functions involved in auditory discrimination (Kiehl et al., 2001). aMMN typically occurs at a latency of 150–200 ms following the presentation of a deviant stimulus. aMMN is observed in the frontal and central electrodes predominantly with polarity reversal at temporal and occipital electrodes (Tada et al., 2020; Escera, 2007).

Only a few aMMN studies on numerical processing exist. However, the brain has been shown to be able to extract numerosity information from sequences of sounds with different frequencies at a pre-attentive level (Ruusuvirta et al., 2007). Furthermore, Ruusuvirta and Astikainen (2016) suggested that automatic numerosity approximation does not rely on attentive individuation mechanisms when it is engaged by small sets of perceptually specific objects. They included standard sequences consisting of three tones, while deviant sequences varied by ±1 tone (two or four tones). The resulting change detection ERP was observed in about 150–200 ms time window and exhibited a fronto-central-parietal topography with a prominent midline component. In line with this, Ruusuvirta and colleagues (2008) demonstrated that changes in small auditory numerosities elicit aMMN responses during passive listening, providing evidence for pre-attentive and automatic encoding of numerosity.

Finally, it is well known that MMN is elicited in response to different changes in auditory and visual features (Näätänen et al., 2007; Stefanics et al., 2014; Male, 2025), but very few studies have examined numerical magnitude changes in auditory or visual domain using MMN (e.g., Hesse et al., 2027; Ruusuvirta & Astikainen, 2016). To date, as far as we know, there are no studies utilizing MMN in numerical and non-numerical magnitude discrimination for both auditory and visual domain in the same participants. Examining both numerical and non-numerical magnitudes as well as auditory and visual domains in the same individuals is important to better understand whether attention allocation is needed for any of these cases, and how similar processing patterns can be found for auditory and visual domains. We wanted to, first, verify that our experimental design indeed elicits auditory and visual MMN responses and, second, to characterize the MMN responses to changes in the number of items presented within the subitizing range. By characterizing the results of the visual and auditory paradigms, we can observe whether the brain detects the changes in the same contrasts of the number of items across modalities.

## Methods

### Participants

Participants were parents of the children followed up longitudinally in the EarlyMath project (ERC grant agreement No. 101002966). They were recruited via maternity clinics and advertisements before the child was born. Data of 126 parents were analyzed for the study.

Twenty-two were excluded for auditory paradigms (bad data, low number of trials) and 21 for the visual paradigms (bad data, low number of trials) for data quality. After pre-processing altogether 113 participants’ data were included in the study, but for some of them only visual or auditory data. The final number of participants for the study was N = 104 (mean age 35.64 years (SD 4.56), range 22.55−45.80 years; 52 females, 52 males) for the auditory paradigms and N = 105 (mean age 35.69 years (SD 4.52), range 22.55−45.80 years; 51 females, 54 males) for the visual paradigms. The exclusion criteria for the research were brain injury, brain tumors, epilepsy, hearing or visual impairment, and skin diseases affecting EEG measurements. In case the participant had medication affecting the central nervous system they were instructed to come to the EEG recording not earlier than three hours after taking their prescribed medication to ensure optimal absorption and minimize interference with neural activity. Before the EEG measurement the hearing threshold, the intensity level (in dB HL) a person can barely detect, was assessed for each participant for frequencies 1000, 500, and 250 Hz in the mentioned order using the standard procedure using Grason-Stadler GSI- 10 audiometer (https://grason-stadler.com/). The study was approved by the ethical committee of the University of Jyväskylä (#1518/13.00.04.00/2021) prior to data collection. Participants gave written informed consent to participate.

### EEG measurement

Electroencephalogram (EEG) was measured in laboratory facilities at the University of Jyväskylä. The continuous EEG was recorded in an acoustically and electrically shielded room using two 32-channel LiveAmp Wireless Amplifiers and BrainVision Recorder 1.24 software (https://www.brainproducts.com/solutions/liveamp/). 64 active EEG electrodes (two actiCAP slim/snap 32-channel sets, sites according to the international 10-10 system) were attached to EasyCap (EasyCap, GmbH, Germany). The EEG was recorded with the sampling rate of 500 Hz using an online band-pass filter from DC to 131 Hz and online referenced to FCz electrode. The ground electrode was located at Fpz. All electrode impedances were aimed to be kept below 20 kΩ. Electrooculogram (EOG) was recorded with two disposable electrodes located at the outer canthi (above the right and below the left outer canthus).

The preparation time for the measurement (including hearing level assessment, EEG electrodes and other sensors) varied between 45−60 min. The total duration of the visit was 2- 3 hours, of which the tasks took approximately 1 hour and 15 minutes.

### Stimuli and procedure

#### Auditory paradigms and stimuli

Three passive auditory EEG paradigms were carried out: a multi-feature paradigm varying in five sinusoidal sound features (Näätänen et al., 2004) and two (deviant-alone) control paradigms. The auditory paradigms utilized six stimuli: 2-part stimulus consisting of two 100 ms long 493.33 Hz sinusoidal tones with 100 ms gap in between the tones presented at 75 dB level as the standard stimulus, and deviant stimuli of 1- part stimulus omitting the second tone from the standard stimulus, 3-part stimulus with additional identical tone inserted with 100 ms gap to the end of the standard stimulus, 2-part stimulus with lower intensity of 65 dB, 2-part stimulus with higher frequency of 554 Hz and 2-part stimulus with shorter gap of 50 ms between two tones. Two control paradigms included either 1-part stimuli or 3-part stimuli presented alone. The stimuli presented in auditory control paradigms were the same as those presented in the multi-feature paradigm. The multi-feature paradigm lasted appr. 25 minutes and was divided into three separate blocks. The auditory control paradigms lasted 2 min and 3 min, respectively.

#### Visual paradigms and stimuli

Three passive visual EEG paradigms were carried out: a multi– feature visual paradigm, with two experimental manipulations, one for numerosity (1–3 dots) and the other for size (large and small) of the dots. Locations of the dots varied pseudorandomly with the following rules: 1) distance of the dots was at minimum two diameters of the medium-sized dot; 2) distance of the dots from the fixation cross was at least the length of diameter of the medium-sized dot; 3) in stimuli with 2 or 3 dots all dots could not all be located on the same visual hemifield; 4) the dots were located within a circular area, whose diameter was 13 cm on the screen. The dot sizes on screen were a large-sized dot 2.5 cm, a medium-sized dot 2 cm, and a small-sized dot 1.2 cm in diameter. Two control paradigms included either stimuli with one dot or stimuli with three dots presented alone. The stimuli presented in visual control paradigms were the same as presented in the multi-feature paradigm. The visual multi-feature paradigm lasted appr. 20 minutes and was divided into five separate blocks. The visual control paradigms lasted appr. 2.5 min each.

#### Auditory stimulus presentation

The auditory stimuli were presented with Presentation® software (www.neurobs.com) and delivered via a loudspeaker located in the ceiling 1 m above the participant. Interstimulus interval (ISI) varied randomly between 650–750 ms in all auditory paradigms. The stimulus intensities were adjusted to 75±0.5 dBA for all the stimuli except 65±0.5 dBA for the attenuated amplitude deviant stimulus (measured with a Brüel & Kjær sound level meter (type 2235). Paradigm order was randomized as well as the order of the three blocks of the multi-feature paradigm. In the auditory multi-feature paradigm, each deviant stimulus was repeated 150 times and a standard stimulus 750 times. In the deviant-alone control paradigm, 1-part deviant and 3-part deviant stimuli were presented alone 150 times.

#### Visual stimulus presentation

The visual stimuli were presented with Presentation® software (www.neurobs.com) and delivered via a screen (Asus VG236H, 23”) located at 1 m from the participant’s eyes. ISI varied randomly between 580–750 ms in all visual paradigms. The stimuli stayed on the screen for 500 ms. Fixation cross was present at the center of the screen continuously. The illuminances (measured with Amprobe LM-120 light meters) of the stimuli were 38-40 Lux (10 cm from screen) and 4.5-5 Lux (at participants’ eye location, 1 meter from screen). Stimuli were presented in the multi-feature paradigm (deviant stimulus order pseudorandomized) and in two separate deviant-alone control paradigms. Paradigm order was randomised as well as the order of five blocks of the multi-feature paradigm. In the visual multi-feature paradigm, each deviant stimulus was repeated 125 times and a standard stimulus 500 times. Both visual control paradigms consisted of 125 repetitions of the stimulus.

The auditory and visual paradigms were counterbalanced between participants. In the visual tasks, participants focused on the central fixation points and were instructed to ignore the series of dots flashed on the screen. During the auditory tasks, they passively listened to a set of tones while watching a silent movie and were instructed not to pay attention to the sounds.

### Data pre-processing

The data were pre-processed with MNE python (Spyder/Python 3.10.4/ iPython 7.32.0). The EEG data were digitally offline filtered with a low cutoff filter of 0.1 Hz and a high cutoff filter of 30 Hz. Channels with multiple artefacts throughout the data were set to “bad” (1.1 bad channels on average per participant) and interpolated before the averaging. EEG epochs with the voltage deflections exceeding ± 150 μV were also excluded before the averaging.

Artefacts related to eye movements and blinks in the data were corrected using ICA (fastICA) in auditory paradigms, but for visual paradigms epochs containing eye movements and blinks were removed. ERPs to each stimulus type were obtained separately by averaging EEG epochs from 50 ms before to 1000 ms after the stimulus presentation and baselined to - 50−0 ms.

### Statistical analysis

The statistical analyses to compare the responses to the deviant stimuli against responses to standard stimuli, as well as numerical deviants against the responses to the same stimuli presented in control paradigms, were done using Threshold-free cluster enhancement (TFCE) method in MNE Python (Mensen & Khatami, 2013). First, the responses to the auditory non-numerical (frequency, gap length, amplitude) and numerical magnitude deviants 1- and 3-part tones were compared with the responses to the 2-part standard stimulus. Second, the responses to the auditory numerical magnitude deviants (1- and 3-part tones) were compared with the responses to the same stimulus presented in auditory control paradigms alone. Third, responses to visual non-numerical (size; large and small) and numerical magnitude deviants (1 and 3 dots) were compared with the responses to the 2-dot standard stimulus. Fourth, responses to visual numerical (1 and 3 dots) magnitude deviants were compared with the responses to the same stimulus presented in visual control paradigms alone. TFCE analysis was carried out for the time window 100−1000 ms. To deal with multiple comparisons, an alpha value (*p*-threshold) of 0.01 was used (*α* 0.05 divided by the number of deviant stimuli used in the auditory multi-feature paradigm that had the most comparisons; this kept the sensitivity of the statistical comparisons comparable across paradigms).

## Results

The ERP waveforms to the standard and deviant stimuli for both the auditory and visual oddball and control experiments followed the expected pattern from previous studies (Figures 1 and 2) (Näätänen et al., 2007; Pazo-Alvarez et al., 2003). The responses to the standard and deviant stimuli in most cases showed clear differences for which the statistical comparisons are described next in addition to the planned comparisons with the responses from the control paradigms.

**Figure 1.**
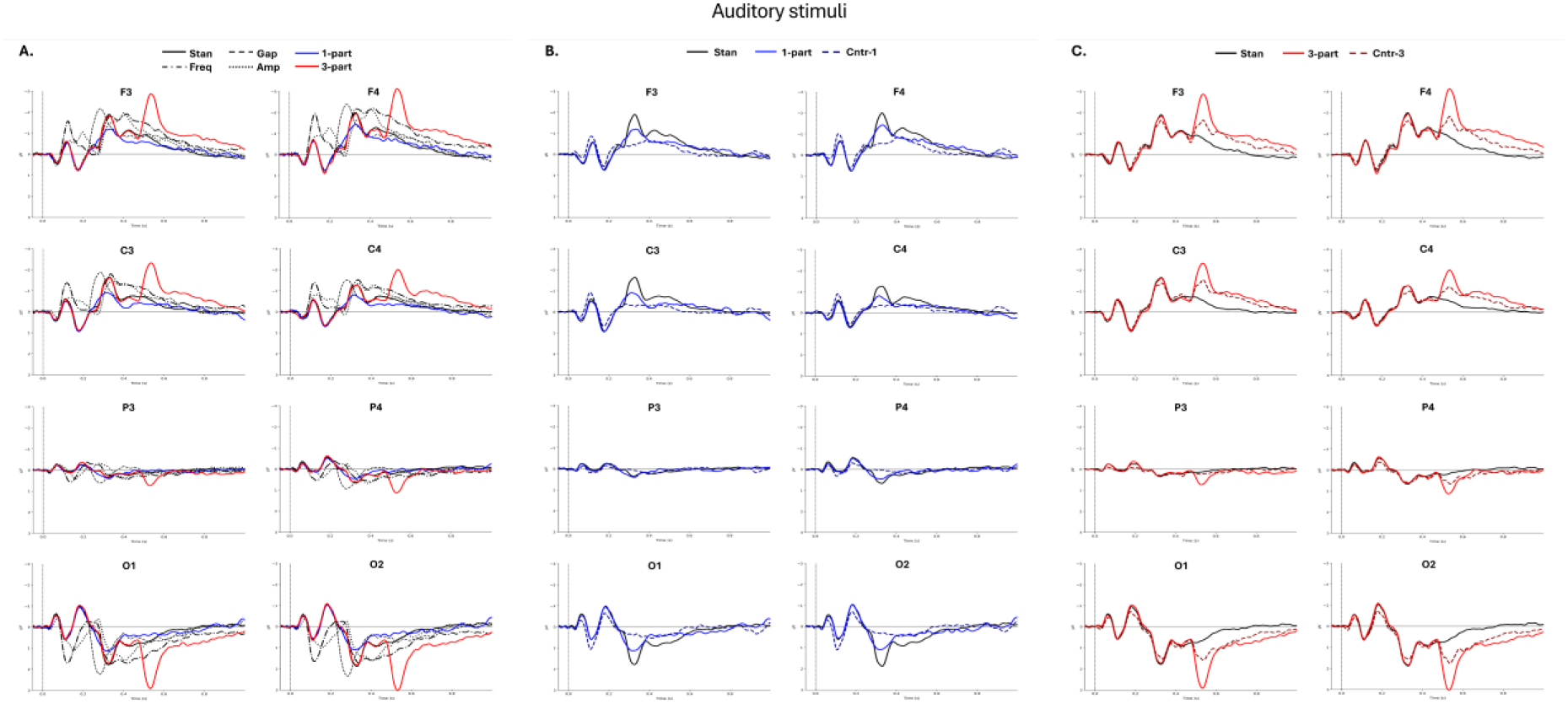
Grand average ERP waveforms (N = 104) for A) all stimuli presented in the auditory multi-feature paradigm, B) for the standard stimuli, 1-part deviant, and 1-part control stimuli presented alone in the control paradigm, and C) for the standard stimuli, 3-part deviant, and 3-part control stimuli presented alone in the control paradigm (negativity up).

**Figure 2.**
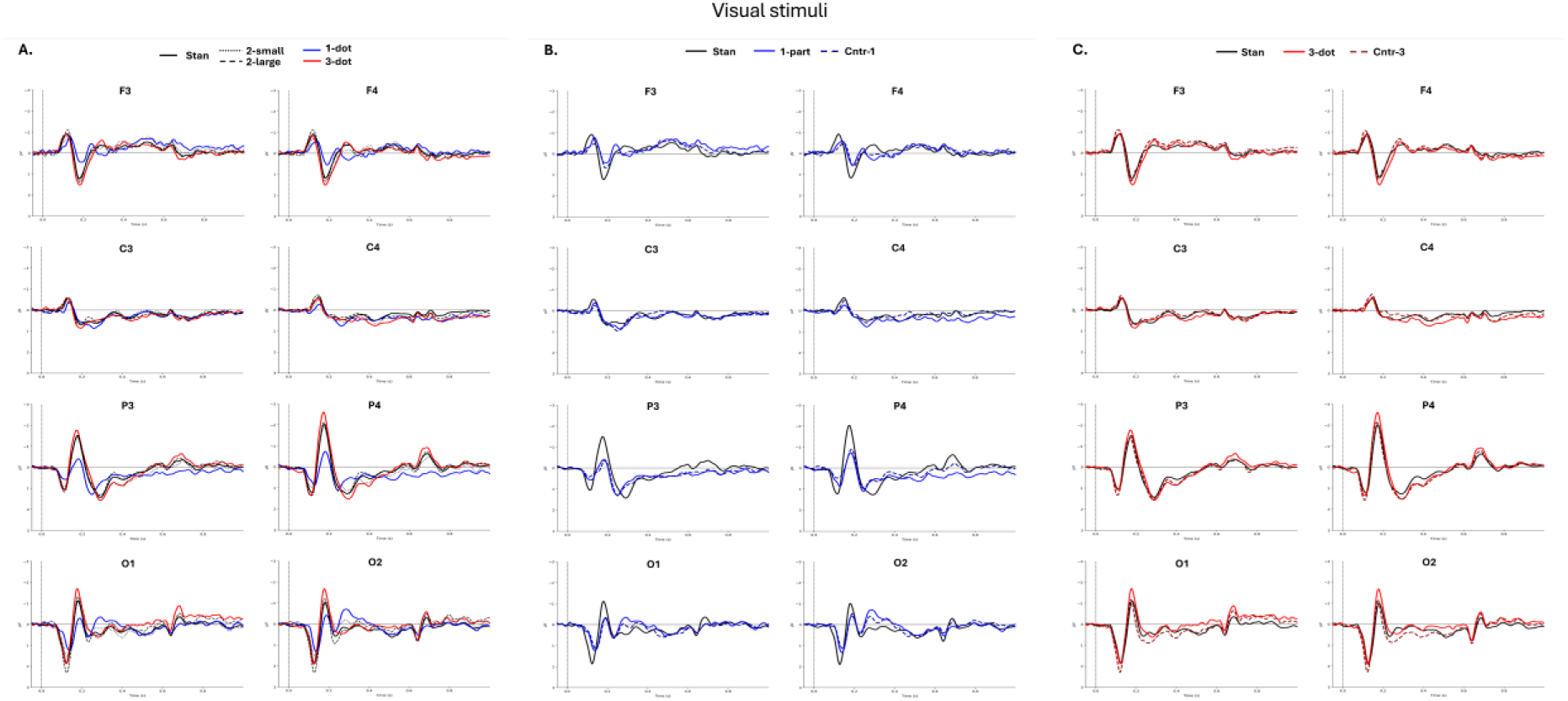
Grand average ERP waveforms (N = 105) for A) all stimuli presented in the visual multi-feature paradigm, B) for the standard stimuli and 1-dot deviant, and 1-dot control stimuli presented alone in the control paradigm, and C) for the standard stimuli, 3-dot deviant, and 3-dot control stimuli presented alone in the control paradigm (negativity up).

### Auditory non-numerical magnitudes

The responses to the deviant stimulus with higher frequency and the 2-part standard stimulus differed from each other within time window of approx. 100−220 ms with the clusters covering frontocentral channels with polarity reversal below the sylvian fissure (see Table 1, see Figure 1A, Figure 3A). Further, there were also two late time window clusters at approx. 350−1000 ms with frontocentral topography and polarity reversal at the parieto-occipital channels.

**Figure 3.**
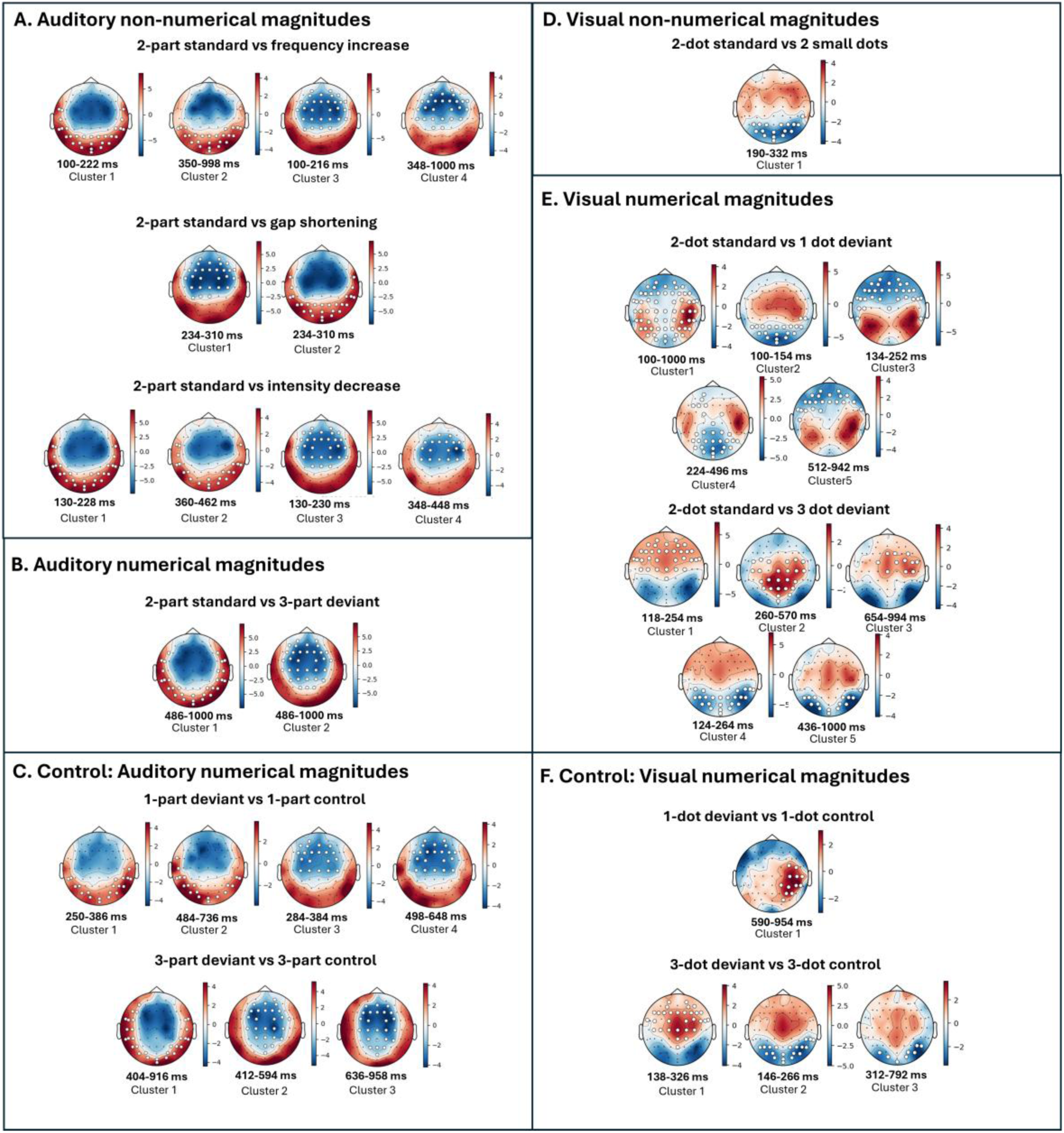
The average t-maps displaying the average *t*-value distribution for TFCE analysis comparisons for auditory (A-C) and visual (D-F) deviant-standard (A, B, D, E) and deviant- control responses (C, F). Sensors forming the clusters are marked by white dots.

**Table 1.**
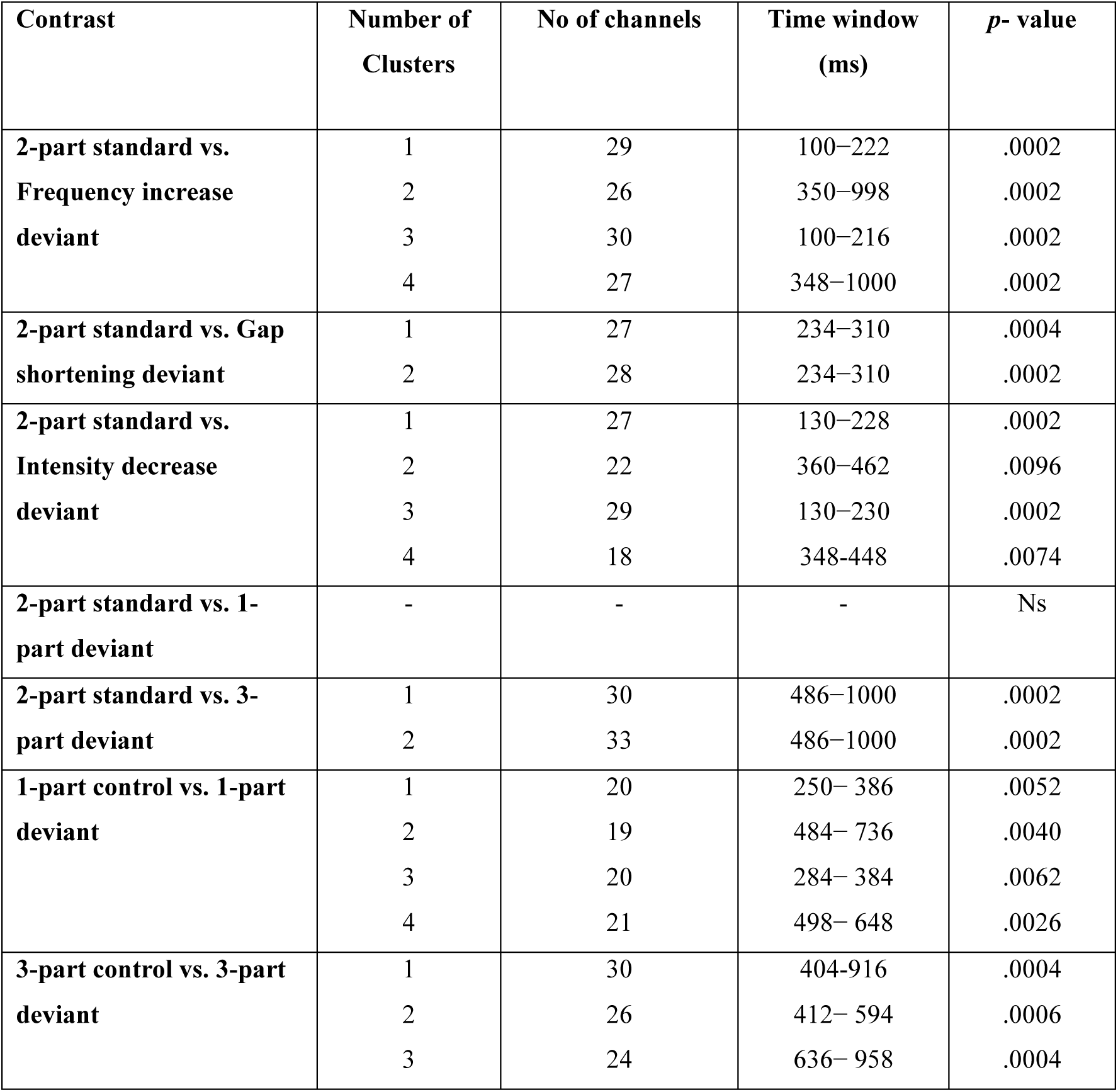
Details of the threshold-free cluster enhancement (TFCE) permutation comparisons between the responses to the auditory stimuli.

| Contrast | Number of Clusters | No of channels | Time window (ms) | *p*- value |
| --- | --- | --- | --- | --- |
| 2-part standard vs. Frequency increase deviant | 1 | 29 | 100–222 | .0002 |
|  | 2 | 26 | 350–998 | .0002 |
|  | 3 | 30 | 100–216 | .0002 |
|  | 4 | 27 | 348–1000 | .0002 |
| 2-part standard vs. Gap shortening deviant | 1 | 27 | 234–310 | .0004 |
|  | 2 | 28 | 234–310 | .0002 |
| 2-part standard vs. Intensity decrease deviant | 1 | 27 | 130–228 | .0002 |
|  | 2 | 22 | 360–462 | .0096 |
|  | 3 | 29 | 130–230 | .0002 |
|  | 4 | 18 | 348–448 | .0074 |
| 2-part standard vs. 1-part deviant | - | - | - | Ns |
| 2-part standard vs. 3-part deviant | 1 | 30 | 486–1000 | .0002 |
|  | 2 | 33 | 486–1000 | .0002 |
| 1-part control vs. 1-part deviant | 1 | 20 | 250– 386 | .0052 |
|  | 2 | 19 | 484– 736 | .0040 |
|  | 3 | 20 | 284– 384 | .0062 |
|  | 4 | 21 | 498– 648 | .0026 |
| 3-part control vs. 3-part deviant | 1 | 30 | 404–916 | .0004 |
|  | 2 | 26 | 412– 594 | .0006 |
|  | 3 | 24 | 636– 958 | .0004 |

The response to the deviant stimulus with shorter gap showed a significant difference to the response to the standard stimulus at approx. 230−310 ms (see Table 1) with frontocentral topography and its reversal at the temporal and occipital areas (see Figure 3A).

The responses to the deviant stimulus with lower amplitude showed significant difference to the responses to the standard stimulus at approx. 130−230 ms and 350−460 ms (see Table 1) both having a frontocentral topography and its reversal at the temporal and occipital areas (see Figure 3A).

### Auditory numerical magnitudes

Responses to the 1-part deviant stimuli and the 2-part standard stimuli did not differ from each other using the alpha level of 0.01. However, the cluster *p*-value was close to the alpha level (*p* < .0195).

Responses to the 3-part stimulus and the 2-part standard stimuli differed statistically significantly in the time window of ca. 485−1000 ms (see Table 1) with frontocentral topography and reversal of polarity at the temporal and occipital channels (see Figure 3B).

### Auditory numerical control

Comparison of the 1-part deviant stimuli and the same stimuli presented alone in the control paradigm showed significant differences in the time windows of 250−380 ms and 480−740 ms (see Table 1) with frontocentral topographies and reversal of polarity in the temporal and occipital temporal areas (see Figures 1B and 3C).

Comparison of the 3-part deviant stimuli and the same stimuli presented alone in the control paradigm showed a significant difference in the time window of 400−960 ms (see Table 1) with frontal, central and parietal topographies that reversed in polarity in the temporal and occipital temporal areas (see Figure 1C and 3C).

### Visual non-numerical magnitudes

Responses to the deviant stimuli with two small dots and the standard stimuli differed from each other at approx. 190−330 ms time window (see Table 2) with occipital scalp distribution as shown in Figures 2A and 3D.

**Table 2.**
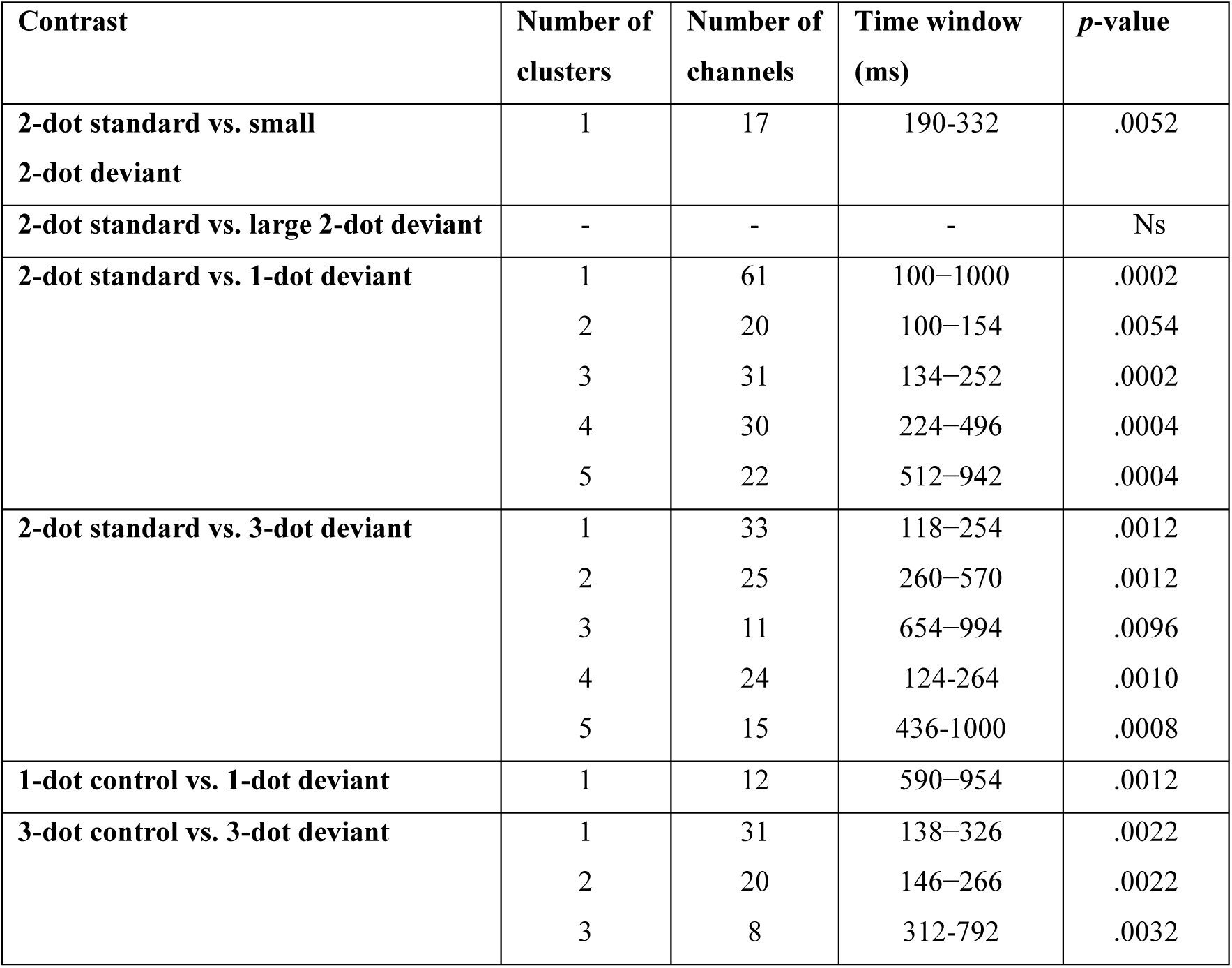
Details of Threshold-free cluster enhancement (TFCE) permutation comparisons between the responses to the visual stimuli.

Responses to the deviant stimuli with two large dots and the standard stimuli showed no statistically significant differences.

### Visual numerical magnitudes

Responses to the 1-dot deviant stimuli and the 2-dot standard stimuli differed from each other at approx. 100−1000 ms time window (see Table 2) with several topographies covering P1, N1 and later time windows with occipitotemporal, occipital, frontal, and parieto-occipital distributions as shown in Figure 3E.

Responses to the 3-dot deviant stimuli and the 2-dot standard stimuli differed from each other at approx. 120−1000 ms time window (see Table 2) with frontocentral and central occipital scalp distribution as shown in Figure 3E.

### Visual numerical control

Responses to the 1-dot deviant stimuli and the same stimuli presented alone in the control paradigm showed a significant difference in the time window of approx. 590−950 ms (see Table 2) with right temporal topography (see Figures 2C and 3F).

Comparison of the 3-dot deviant stimuli and the same stimuli presented alone in the control paradigm showed a significant difference in the time window of approx. 140−800 ms (see Table 2) with frontocentral and bilateral occipital topography (see Figures 2C and 3F).

## Discussion

The aim of this study was to understand how the adult brain passively responds to changes in numerical and non-numerical magnitudes in the auditory and visual domains. Here, numerical and non-numerical magnitudes refer to non-symbolic representations of the number of tones, frequency, intensity, and gap length in the stimulus as well as the number and size of dots. We used auditory and visual MMN paradigms, and deviant-alone control paradigms to evaluate neural responses to changes in the number of visual and auditory elements. For the auditory stimuli, differences were observed for each standard-deviant contrast except for the comparison between the responses for the 2-part standard stimulus and the 1-part deviant stimulus responses. However, the responses to both numerical deviants differed from the responses to the same stimuli presented alone in the control paradigm. For the visual stimuli, differences were observed for each standard-deviant contrast except for the comparison between the responses to the 2-dot standard and the deviant stimulus with 2 larger dots. When comparing responses for the numerical deviants to the responses for the same stimuli from the control paradigm, difference was observed only between the responses to the 3-dot stimuli, but not between the 1-dot stimuli. Consistent with previous evidence, the present results indicate that by using the multi-feature oddball paradigm it is possible to effectively elicit neural responses to both non-numerical and numerical changes (Näätänen et al., 2004, 2007; Pakarinen et al., 2009; Li et al., 2025; Widmann et al., 2026).

### Auditory responses

In line with earlier aMMN work (Garrido et al., 2009; Näätänen et al., 2007), the responses to all three non-numerical deviant stimuli showed differences to those of the standard stimuli with similar frontocentral topographies at the difference starting approximately 100 ms after deviancy onset and lasting several hundred milliseconds. When examining the responses to the numerical deviants, responses to the 2-part standard stimuli and the 1-part deviant stimuli did not differ from each other while responses to the 3-part stimulus and the standard stimuli differed starting at 86 ms after deviancy onset with frontocentral topography and reversal of polarity at the temporal and occipital channels. Topographies for all the aMMN responses were very similar by visual inspection suggesting activation of change detection processes in similar brain regions (i.e., generator sources) across the different contrasts.

Brain responses to the 3-part deviant stimuli and the same stimuli presented alone in the control paradigm differed significantly between 400−960 ms (see Table 1). These changes were strongest over the frontocentral area with reversal of polarity in the occipital and temporal areas (see Figure 2). If physically identical stimuli produce different ERP responses depending on context (deviant vs. control), MMN is likely present reflecting predictive auditory processing related to change detection rather than simple sensory processing (Widmann et al., 2025), although it should be noted that the time window was relatively late for a typical MMN response.

Interestingly, no significant difference was observed between the responses to the 2-part standard and the 1-part deviant stimuli at the alpha level of 0.01; however, the cluster-level *p*- value approached significance (*p* < .0195). When comparing the 1-part deviant stimuli and the same stimuli presented alone in the control paradigm, significant difference was observed. These results could indicate the presence of a change detection response combined with the marginally significant results in the standard vs. deviant comparison using Bonferroni correction albeit with a weak effect.

Although both contrasts involved either addition or removal of one sound element (although the stimulus ratio was different) from the standard, discrimination was more robust for the 2- part versus 3-part tones than for the 2-part versus 1-part contrast. The weaker 2-part versus 1- part discrimination may reflect the omission-like nature of the 1-part deviant. Studies of omission demonstrate MMN-like responses to the absence of expected auditory events (Yabe et al., 1997, 1998). The difference in our results and those of Yabe and colleagues (1997, 1998) could be due to shorter ISIs used in their studies and related to the integration of information in shorter intervals into perceptual objects that might be disrupted for longer ISIs. Further, in our study we used a stricter alpha level (due to multiple comparisons) that might have impacted the sensitivity to find a statistically significant effect.

Within the predictive coding framework, the brain continuously generates predictions about forthcoming sensory events, and violations of these predictions elicit prediction-error signals that support the updating of internal models (Friston & Kiebel, 2009; Garrido et al., 2009).

Consequently, the omission of an expected auditory component may generate a weaker prediction-error signal than the addition of an unexpected component, potentially accounting for the larger discrimination effect observed for the 2-part versus 3-part contrast. This interpretation is further supported by evidence indicating that MMN reflects active predictive processing rather than passive sensory adaptation (Bendixen et al., 2009; Wacongne et al., 2011). Similarly, Timm et al. (2011) reported greater neural sensitivity to the presence than to the absence of frequency modulation. Although their paradigm examined feature-based rather than numerosity-based discrimination, their findings nevertheless suggest a broader asymmetry in auditory change detection, whereby the addition of auditory information elicits stronger neural responses than its omission.

Consistent with previous aMMN studies demonstrating that the auditory system can encode small numerosities at both preattentive and attentive levels (Ruusuvirta et al., 2007; Ruusuvirta et al., 2016), our participants showed change detection responses between the 2- and 3-part sound sequences. Although the studies by Ruusuvirta et al. primarily investigated ratio-based numerosity changes rather than the absolute quantity contrasts examined here, both sets of findings suggest that auditory numerical information can be represented automatically. However, the absence of a significant effect for the one- versus two-sound contrast indicates that the discriminability of auditory numerosities may vary across numerical contrasts, with some quantity changes eliciting weaker neural responses than others. The present findings provide evidence that numerical changes in auditory sequences influence early automatic neural predictions which precede conscious numerical judgments.

### Visual responses

We first examined if changes in dot sizes elicit the visual mismatch negativity (vMMN) responses verifying that the stimulation parameters would be sufficient to elicit a vMMN also for the numerical changes. Our results indicate that the deviant containing two small dots compared to the medium sized dots elicited a significant change detection response, characterized by an early occipital negativity between 190–330 ms. This spatiotemporal pattern is consistent with earlier studies which have localized the vMMN to parieto-occipital and occipital cortices (Male, 2025; Pazo-Alvarez et al., 2003). In contrast, with two large dots, the responses to the deviant elicited no differences from the standard stimulus responses. These findings suggest that the visual systems are sensitive to changes in surface, to a certain extent consistent with the ratio of change (ca. 2.8 for smaller versus standard size dots, 1.6 for larger versus standard size dots). The greater sensitivity observed for the 2.8 ratio relative to the 1.6 ratio suggests that discrimination of cumulative area may depend on the proportional difference between magnitudes rather than their absolute values. This interpretation aligns with empirical evidence demonstrating ratio-dependent magnitude discrimination in adults (Buckley & Gillman, 1974). Furthermore, Bonny and Lourenco (2023) reported behavioral and electrophysiological ratio effects during cumulative area judgments, indicating that surface-area representations are sensitive to relative differences between quantities. Alternatively, the stronger response to the 2.8 relative to the 1.6 ratio may reflect greater perceptual salience of surface-area changes arising from continuous visual properties (Gebuis & Reynvoet, 2013; 2012).

Evidence for automatic numerosity processing has also been studied in visual modality (Gennari et al., 2023; Hesse et al., 2017). One visual oddball study demonstrated that unattended changes in small numerosities using dot stimuli elicited a visual mismatch negativity (vMMN), suggesting the existence of preattentive numerical processing (Hesse et al., 2017). Similarly, our work focused on the visual dot arrays within the subitizing range. Responses to the 1-dot deviant stimuli and the 2-dot standard stimuli differed from each other at approx. 100−1000 ms time window (see Table 2) with several topographies covering P1, N1 and later time windows with occipitotemporal, occipital, frontal, and parieto-occipital distributions. Responses to the 3-dot deviant stimuli and the standard stimuli differed from each other at approx. 120−540 ms time window (see Table 2) with frontocentral and central occipital scalp distributions. The 3-dot deviant stimuli produced longer responses than the 1- dot deviant stimuli, suggesting more complex neural engagement. In line with the Hesse et al., 2017 study, the onset of the MMN is roughly in line with our results. However, we showed a longer lasting differential response compared to Hesse et al. 2017. The topography of the MMN was similar between the studies with stimulus contrasts showing differences at the parieto-occipital sites. However, it should be noted that there were several methodological differences in the experimental design between the studies that might have impacted the duration of the MMN response.

When compared with the control paradigm, the 3-dot deviant stimuli elicited a genuine change detection response as reflected in the early occipital effect 130−330 ms. Similarly, responses to the 1-dot deviant showed a significant difference when compared to those of the 2-dot standard and also differed when compared to the responses to the same stimuli presented in the control paradigm. This suggests that both numerical contrasts trigger a change detection response.

### Limitations of the study and future directions

There are some limitations that need to be taken into account. First, due to time constraints, we were not able to run a full set of control experiments. Optimally, the control experiments would have included an equal probability condition where the deviant stimuli would be presented in a non-oddball sequence with the same probability as in the oddball condition. This would have shown stronger evidence for specific change detection processing.

Additional control experiments could have examined the effects of stimulus features covarying with numerosity such as visual density or auditory duration. However, our main purpose was not to examine whether numerical processing is independent of other sensory cues present in the stimuli, but rather to present evidence that changing the number of items in the stimuli can elicit change detection responses regardless of the modality.

Future studies could therefore include more control experiments as well as expand the range of numerosities examined beyond the subitizing range. This would allow conclusions on how similar or different the processing for small and large numerosities are given the theoretical prediction on the separate mechanisms for these (Ansari et al., 2007; Hyde & Spelke, 2009; Hyde, 2011; Nieder, 2025).

### Conclusions

We conducted auditory and visual experiments to investigate mismatch negativity (MMN) responses in adults as an index of pre-attentive processing of different numerical (small numerosities 1–3) and non-numerical magnitudes. Overall, we observed positive evidence for both auditory and visual MMN. Auditory and visual MMN were elicited for changes in numerical magnitude but more robustly to contrast 2 vs. 3 than to contrasts 2 vs. 1. While our findings suggest that numerical processing was measured to some extent, the precise nature of the change detection processing between one and two non-symbolic numerosities at the pre-attentive level remains unclear and processing of quantity 1 versus 2–3 might rely on partially distinct processes.

The presence of both auditory and visual MMN responses suggests that changes in non-symbolic small numerosity can be detected without conscious attention, which is consistent with earlier results of pre-attentive numerical processing (Hesse et al., 2017; Ruusuvirta & Astikainen, 2016). Furthermore, an earlier EEG study using auditory and visual stimulation was able to decode numerical processing across modalities (Gennari et al., 2023). Together with the MMN responses in our study across both auditory and visual modalities the result could be interpreted to support the existence of abstract representations of magnitudes (Dehaene et al., 1998).

## Acknowledgments

We would like to thank the research assistants for data collection and all the participants for taking part in the study.

The project was funded by the European Union (ERC, EARLYMATH, 101002966). Views and opinions expressed are however those of the author(s) only and do not necessarily reflect those of the European Union or the European Research Council. Neither the European Union nor the granting authority can be held responsible for them. SM was funded by the Finnish National Agency for Education.

## Notes

### Competing Interest Statement

The authors have declared no competing interest.

